# Spartin-mediated lipid transfer facilitates lipid droplet turnover

**DOI:** 10.1101/2023.11.29.569220

**Authors:** Neng Wan, Zhouping Hong, Matthew A. H. Parson, Justin Korfhage, John E. Burke, Thomas J. Melia, Karin M. Reinisch

**Affiliations:** Department of Cell Biology, Yale University School of Medicine, New Haven, CT 06520, USA; Department of Biochemistry and Microbiology, University of Victoria, Victoria, BC, Canada V8W2Y2; Department of Biochemistry and Molecular Biology, University of British Columbia, Vancouver, BC, Canada V6T 1Z3

## Abstract

Lipid droplets (LDs) are organelles critical for energy storage and membrane lipid homeostasis, whose number and size are carefully regulated in response to cellular conditions. The molecular mechanisms underlying lipid droplet biogenesis and degradation, however, are not well understood. The Troyer syndrome protein spartin (SPG20) supports LD delivery to autophagosomes for turnover via lipophagy. Here, we characterize spartin as a lipid transfer protein whose transfer ability is required for LD degradation. Spartin co-purifies with phospholipids and neutral lipids from cells and transfers phospholipids in vitro via its senescence domain. A senescence domain truncation that impairs lipid transfer in vitro also impairs LD turnover in cells while not affecting spartin association with either LDs or autophagosomes, supporting that spartin’s lipid transfer ability is physiologically relevant. Our data indicate a role for spartin-mediated lipid transfer in LD turnover.

**Significance:** The Troyer syndrome protein spartin was proposed to function as a lipophagy receptor that delivers lipid droplets, organelles key for energy storage and membrane lipid homeostasis, to autophagosomes for degradation. We identify an additional function for spartin as a lipid transfer protein and show its transfer ability is required for lipid droplet degradation, including by lipophagy. Our data support that protein-mediated lipid transfer plays a role in lipid droplet turnover. Moreover, in spartin’s senescence domain we have discovered a new lipid transport module that likely also features in still undiscovered aspects of lipid droplet biology and membrane homeostasis.

## Introduction

Comprising a core of neutral lipids surrounded by a monolayer of glycerophospholipids, lipid droplets (LDs) are storage organelles for lipids prior to their mobilization as precursors in membrane lipid biosynthesis or for metabolic energy. As such these organelles play key roles in physiology and metabolism. The mechanisms by which lipids are transferred to or mobilized from LDs remain poorly understood but have been speculated to involve protein-mediated lipid transfer, presumably at sites of close apposition between LDs and other organelles (1, 2). Thus, the lipid transport ability of mitoguardin-2, between mitochondria and LDs, plays a role in LD biogenesis (3) ; and lipid transport proteins (LTPs) in the VPS13-family and the VPS13-like protein ATG2 localize to LDs (4, 5), although their function in LD biology is not well established. The protein spartin (SPG20) also localizes to LDs and participates in their turnover (6), and it is recently proposed as a receptor that delivers LDs to autophagosomes for degradation via macrolipophagy (7). Here we report that spartin additionally has the ability to bind and transfer lipids in vitro and that impairment of lipid transport abrogates its function in LD degradation.

Spartin is present in higher eukaryotes, and mutations in the human gene are causative of the rare neuronal disease Troyer’s syndrome (8), a form of hereditary spastic paraplegia. Spartin is predicted to comprise a microtubule interacting (MIT) domain, a pleckstrin homology (PH)-like domain, and a “senescence domain” at the C-terminus (Fig 1A). Additionally, spartin-like proteins, with in-tandem PH-like and senescence domains but lacking the MIT domain, are present in plants and certain fungi. In human spartin, a motif upstream of the PH-like domain binds to LC3A/C proteins present on the forming autophagosome, and two amphipathic helices in the senescence domain (helices 1 and 2) mediate association with LDs (7). The senescence domain is predicted to comprise four helical segments,and has not been extensively characterized structurally or functionally. We show that spartin’s ability to bind and transfer lipids resides in the senescence domain. Moreover, a truncation in this domain that impairs lipid transfer in vitro also impairs LD degradation in cells even as spartin localization to LDs or LC3-positive autophagosomes is not affected. Our findings indicate the senescence domain as a novel lipid transfer module and demonstrate human spartin as a lipid transport protein involved in LD turnover.

**Figure 1:**
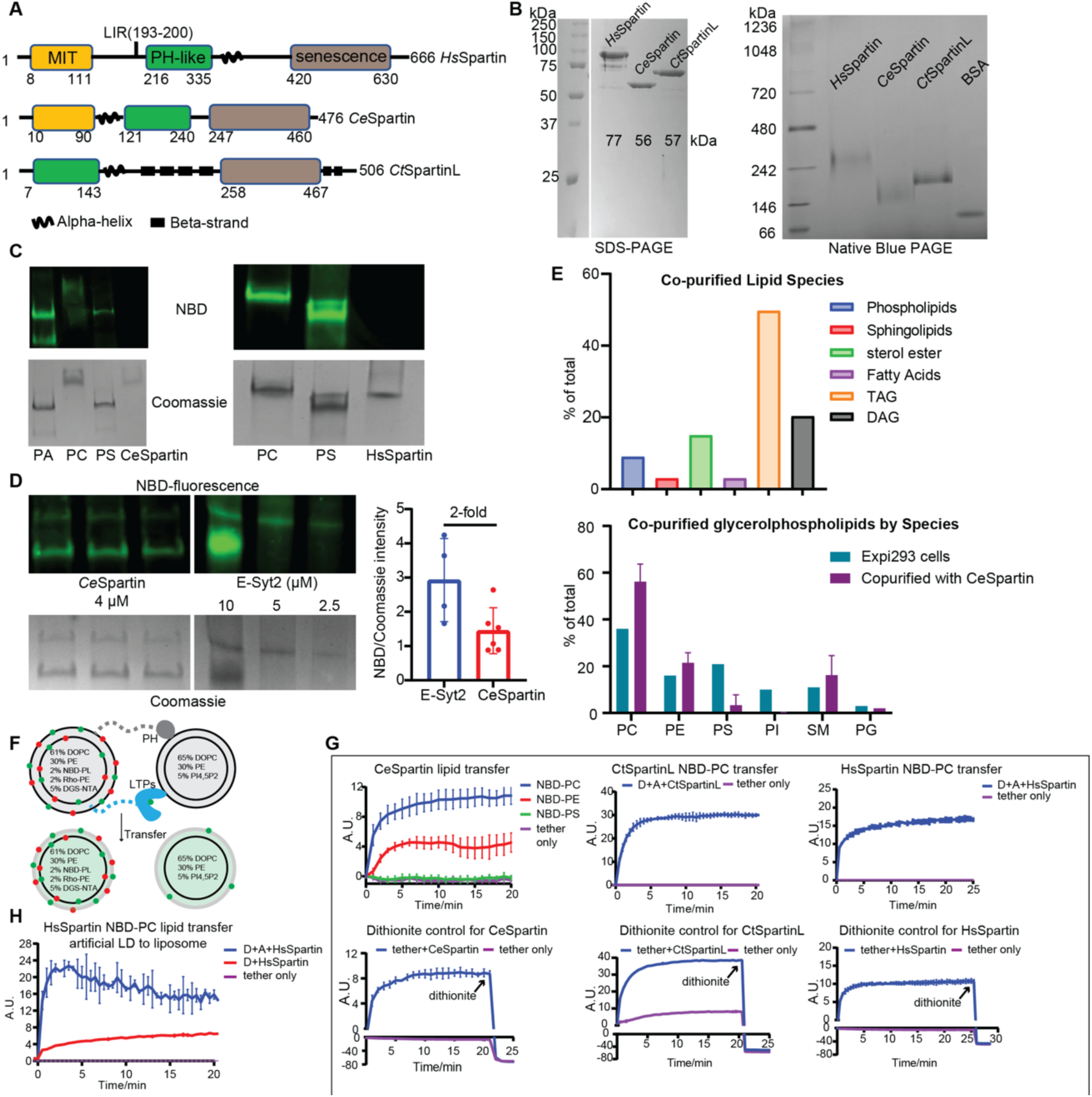
Spartin proteins bind and transfer lipids. **A**: Domain diagrams of spartin proteins including *Hs*Spartin, *Ce*Spartin and *Ct*SpartinL. *Hs*Spartin and *Ce*Spartin have three domains: microtubule-interacting (MIT) domain, PH-like domain and senescence domain. *Hs*Spartin has a LC3A interacting region (LIR: 193-200) (7). Spartin-like protein from Chaetomium thermophilum (*Ct*SpartinL) lacks the MIT domain. **B:** Left: SDS-PAGE of purified spartin. Right: Native blue PAGE analysis of same spartin proteins indicates that all three proteins likely trimerize. **C**: In vitro lipid binding assays show that *Ce*Spartin and *Hs*Spartin bind lipids. *Ce*Spartin and *Hs*Spartin were incubated with NBD-phospholipids or solvent (no lipids) and analyzed by native gel electrophoresis, where migration depends on charge-mass ratio of the protein-lipid complexes. **D**: Quantitative in vitro lipid binding assay estimates *Ce*Spartin binds one lipid/protein molecule. CeSpartin (4μM) and E-Syt2 (10, 5.0, 2.5 μM) were incubated with NBD-PS and analyzed as in C. (At 10 μM, E-Syt2 migrates as two species). The Coomassie density allows determination of the number of proteins in the sample, whereas the NBD fluorescence indicates quantity of lipids bound by this number of proteins. Because each Extended Synaptotagmin monomer is known to bind 2 lipids, we can extrapolate how many lipids the fluorescence signal associated with spartin represents. **E**: Lipidomic MS analysis of *Ce*Spartin purified from Expi293 cell shows that it binds neutral lipids (sterol esters, TAG and DAG) and glycerophospholipids. Percentage of co-purified phospholipids is plotted against phospholipids profile of Expi293 cells. **F**: Schematic of in vitro FRET-based lipid transfer assay. **G**: Lipid transfer assays show that spartin proteins transfer lipids between liposomes. NBD-PC/PE/PS substrates were tested for *Ce*Spartin, while only NBD-PC transfer assays were performed for *Hs*Spartin and *Ct*SpartinL. Proteins were purified from Expi293 cells and used at 0.25 μM (1:800 protein:lipid ratio). Each experiment was performed in triplicate; SD is indicated. In control experiments, to verify that the increase in fluorescence is not due to fusion of the donor and acceptor liposomes, we added dithionite following the assay. Dithionite reduces solvent exposed NBD and quenches its fluorescence but does not affect NBD-lipids in the liposome lumen. After dithionite addition, NBD-fluorescence is reduced equally whether spartin is present or not, indicating that only the outer leaflet of the liposomes changed in the course of the lipid transfer reaction, not both leaflets as would be expected had fusion taken place (13). **H**: In vitro lipid transfer assay shows that *Hs*Spartin (0.1 μM, 1:800 protein:lipid ratio) robustly transfers NBD-PC from artificial LDs to liposomes (n=3). Artificial LDs prepared as in (14) were used as lipid donors instead of liposomes. See Fig. S1A for characterization of the artificial LDs by positive stain electron microscopy.

## Results and Discussion

### Spartin binds and transfers lipids

That many lipid transfer proteins were first identified as tethers between organelles and only subsequently shown to transfer lipids (9, 10) and that spartin appears to function as a tether between LDs and the autophagosome during LD turnover (7) raised the possibility that it might also have lipid transfer ability. As a first assessment of this notion, we asked whether, like known lipid transporters, spartin can bind lipids. For these experiments we isolated full-length versions of spartin from humans or *C. elegans* (HsSpartin, CeSpartin) or a spartin-like protein from the fungus *Chaetomium thermophilum* (CtSpartinL). We chose to work with CeSpartin and CtSpartinL in addition to HsSpartin because their yields of purified protein were higher and so more suitable for biochemistry. All three proteins were expressed in Expi293 cells or alternatively in bacteria (for CeSpartin and CtSpartinL) and purified via an N-terminal affinity tag (3XFLAG tag for constructs made in Expi293 cells and a hexahistidine tag for those made in bacteria), and the affinity tag was removed via proteolytic cleavage. All three proteins multimerize as assessed by native blue gel, their sizes consistent with trimerization (Fig. 1B). When we incubated the purified proteins with NBD-labeled fluorescent glycerophospholipids, they co-migrated with fluorescence on native gels, indicating that they bind these lipids (Fig. 1C). By comparing the fluorescence that co-migrated with spartin versus that co-migrating with a well characterized lipid transport protein (E-Syt2, (11)), we estimate that one lipid binds per spartin molecule (Fig. 1D).

To discover which lipids spartin might bind in vivo, we purified CeSpartin from mammalian cells (Expi293 cells), washing FLAG-resin immobilized CeSpartin extensively (with 4 column volumes over 1 hour) to remove as much as possible non-specifically bound lipids, and analyzed spartin-associated lipids using liquid chromatography/liquid chromatography/mass spectrometry (LC/LC/MS). Consistent with a role in LD biology, we found that CeSpartin associated with lipids normally enriched in LDs (Fig. 1E). These were predominantly glycerolipids and sterol esters, the neutral lipids in the LD core, as well as phosphatidylcholine (PC) and phosphatidylethanolamine (PE), the major glycerophospholipids in the surrounding monolayer. PC and PE were enriched as compared to their abundance in cells, suggesting a preference as compared to other glycerophospholipids (phosphatidylserine or phosphatidylinositol).

Using a well-established FRET-based assay (4), we next assessed whether spartin or spartinL can transfer lipids between membranes in vitro (Fig 1F). In the assay, the candidate LTP is tethered between donor and acceptor liposomes, mimicking localization to sites of organellar apposition. Lipid transfer between membranes and the LTP is stochastic and rate determining in the transfer reaction (9), and tethering assures that the LTP associates with liposomes sufficiently for the transfer to occur (discussed in (3)). We tethered together donor and acceptor liposomes using a previously described linker construct (12). The linker has an N-terminal hexahistidine tag allowing for binding to Ni-NTA lipids (5%) in the donor liposome and a C-terminal PH domain allowing for binding to phosphoinositide PI(4,5)P_2_ (5%) in the acceptor liposomes. Spartin or SpartinL constructs, purified from Expi293 cells via affinity purification as before, feature a C-terminal linker fused to a hexahistidine sequence, allowing tethering to the donor liposomes.Donor liposomes initially contain both Rhodamine (Rh)-PE (2%) and a nitrobenzoadiazole (NBD)-labeled lipid (2%), so that FRET between lipid-associated Rh and NBD quenches the NBD fluorescence. Acceptor liposomes initially lack fluorescent lipids. Addition of an LTP and consequent transfer of fluorescent lipid species from donor to acceptor liposomes results in their dilution, decreased FRET between Rh and NBD, and increased NBD fluorescence. Using NBD-PC, we find that addition of CeSpartin, HsSpartin or CtSpartinL to the liposomes all result in increased NBD fluorescence, consistent with an ability to transfer glycerophospholipids (Fig. 1G). Moreover, in further experiments with CeSpartin, we observed fluorescence increases consistent with NBD-PC or NBD-PE transfer, although NBD-PE transfer was less robust, but not NBD-PS transfer, indicating selectivity for the glycerophospholipids enriched in LDs. We used a dithionite quenching assay (13) to exclude the possibility that the observed fluorescence increases are due to spartin-mediated fusion of donor and acceptor liposomes (Fig. 1G).

Because Spartin localizes to LDs in cells (6), we further asked whether it can transfer lipids between LDs and other membranes. We used the same FRET-based assay as above, except that we prepared artificial LDs (14) instead of liposomes as lipid donors. We find that HsSpartin robustly transfers NBD-PC from the artificial LD preparation (characterized in Fig S1A) to liposomes (Fig. 1H).

### The senescence domain is required and sufficient for lipid transfer

We carried out lipid transfer experiments with truncation constructs to identify the portion of spartin/spartinL responsible for transfer activity. As noted, CtSpartinL lacks an MIT domain, suggesting that the lipid transfer function resides in the PH-like or/and senescence domains. Consistent with this, a CeSpartin construct lacking the MIT domain (CeSpartin_PH-SD_, residues 121-476) still transfers lipids (Fig. 2A). In contrast, a construct comprising only the PH-like domain (CeSpartin_PH_, residues 95-240, or CtSpartinL_PH_, residues 7-245) does not transfer lipids (Fig. 2A, B), indicating that the senescence domain is required for transfer activity. Further confirming the importance of this domain for lipid transfer, removing either the first two or last two helical segments of CtSpartinL’s senescence domain abrogated lipid transfer activity (CtSpartinL_ΔH1H2_, CtSpartinL_ΔH3H4_), and a mutation in the first helix (A290P, CtSpartinL_A290P_) similarly disrupted both lipid binding and transfer activity (Fig. 2B,C). An analogous mutation in the senescence domain of HsSpartin that occurs in Troyer’s syndrome patients (A442P) (15) also disrupts lipid transfer in vitro, as does deletion of helical segments 3 and 4 of the senescence domain (HsSpartin_ΔH3H4_) (Fig. 2D).

**Figure 2:**
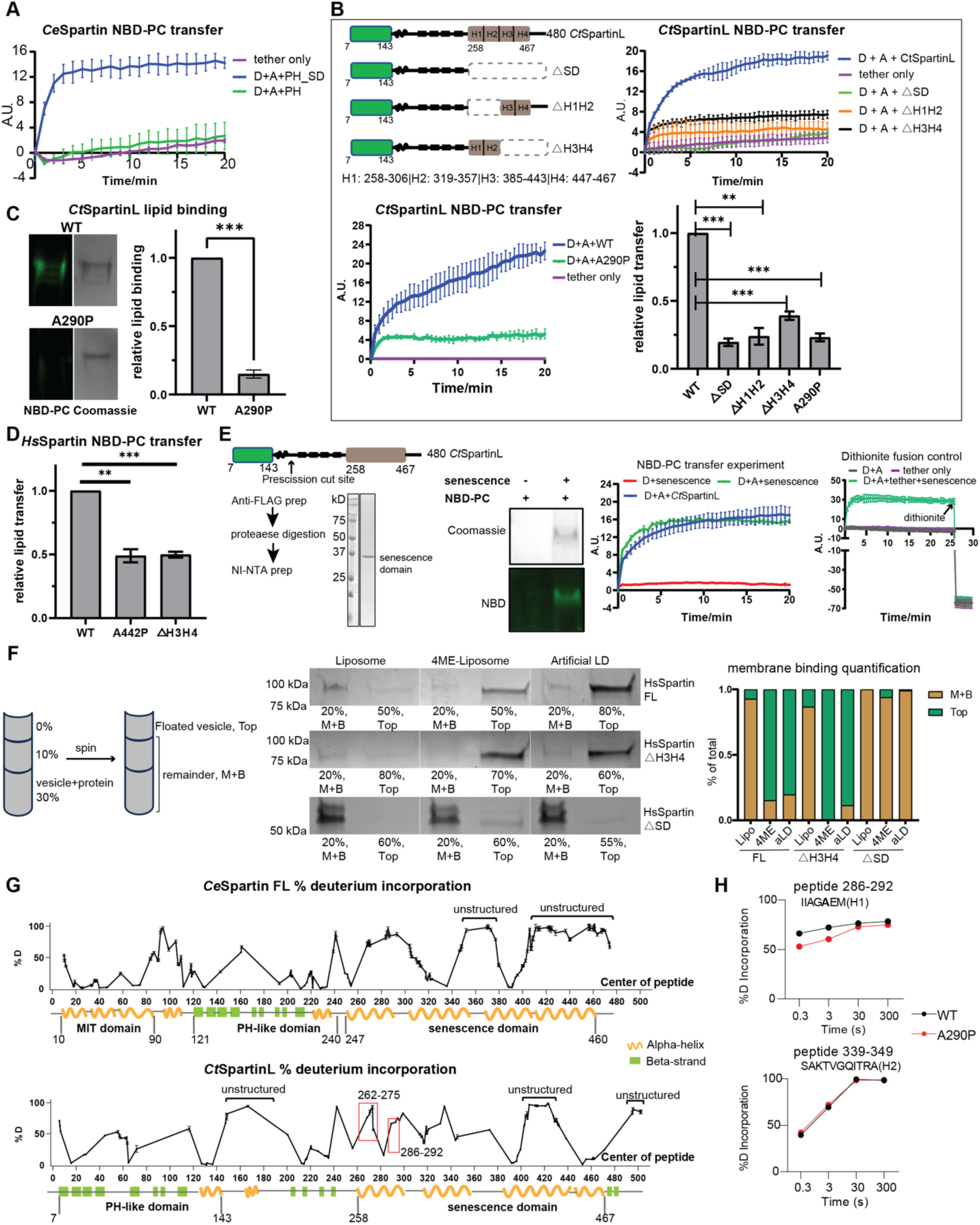
Senescence domain is necessary and sufficient for lipid transfer. **A**: Lipid transfer activity of *Ce*Spartin depends on its senescence domain (SD). A PH+SD construct of *Ce*Spartin (aa 121-476) retains lipid transfer activity, while the PH domain alone (aa 95-240) is transfer incompetent. n=3; SD indicated. **B**: Lipid transfer by *Ct*SpartinL requires intact senescence domain. Constructs are indicated at upper left: either the entire (ΔSD) or half of the senescence domain (ΔH1H2 or ΔH3H4 of *Ct*SpartinL) was deleted. All three truncated constructs had significantly impaired lipid transfer activity (upper right). A point mutation (A290P) within H1 also disrupted lipid transfer by *Ct*SpartinL (lower left). Experiment was performed in triplicate, with SD indicated. Lower right: quantification of lipid transfer 20 minutes after addition of either mutant constructs or WT protein. **C**: The *Ct*SpartinL_A290P_ mutant has decreased lipid binding ability. Lipid binding assay for WT *Ct*SpartinL and *Ct*SpartinL _A290P_ performed as described before. *Ct*SpartinL_A290P_ had only ∼15% lipid binding capacity of WT protein (n=3). **D**: Lipid transfer activity of HsSpartin requires intact senescence domain. NBD-PC transfer experiments were performed for HsSpartin_ΔH3H4_ and HsSpartin_A442P_, along with WT *Hs*Spartin. Both mutants have reduced lipid transfer ability (quantitation shows the end point of lipid transfer experiment). n=3; SD is indicated. **E**: Left panel: Strategy for production of a senescence-domain-only construct from *Ct*SpartinL. 3XFLAG-(*Ct*SpartinL 7-480)_Precission_-6HIS construct was engineered with a Prescission protease site (arrow). The construct was isolated first using anti-FLAG resin, digested with protease, and further purified using the C-terminal 6HIS tag to capture the senescence domain fragment. Middle panel: senescence domain fragment co-migrates with NBD-PC on native gels. Right panel: NBD-PC transfer assay shows that the lipid transfer activity of the senescence domain fragment (80 μM lipid, 0.1 μM protein) is comparable to the full-length construct’s (CtSpartinL, 200 μM lipid, 0.25 μM protein); n=3 for all curves. **F:** In vitro flotation assay shows that HsSpartin binds directly to artificial lipid droplets and liposomes with membrane packing defects. Purified proteins were incubated with liposomes, liposomes comprising 4-ME lipids, or artificial LDs at RT for 30min, then analyzed by density gradient ultracentrifugation (in Optiprep). Post ultracentrifugation, floated vesicles or LDs on the top and the remainder (Middle+Bottom, M+B) were collected for SDS-PAGE analysis (% of M+B or top fractions loaded onto gels is indicated). Quantification on the right shows that HsSpartin_△H3H4_ construct binds to membranes (containing 4ME-lipo) and artificial LDs like WT HsSpartin, whereas HsSpartin_△senescence_ construct does not bind; n=2. **G**: Percentage of deuterium incorporation for *Ce*Spartin (purified from *E. coli*, after 3s at 20°C) and *Ct*SpartinL (purified from Expi293 cells, after 3s on ice) after deuterium exposure. Each data point represents the central residue of a single peptide; data shown were corrected using a fully deuterated control. Predicted secondary structure is indicated. Experiments were performed in triplicate; SD indicated. Extensive unstructured regions (highest level of deuterium incorporation) are labeled. For CtSpartinL, red rectangles highlight regions with significant differences in deuterium incorporation in CtSpartinLA290P versus WT (peptides: 262-275, 286-292). **H**: *Ct*SpartinL A290P shows significant differences of deuterium incorporation compared with *Ct*SpartinL WT within H1. Upper panel: percentages of deuterium incorporation for the peptide 286-292, where changes were observed at different exposure times. In contrast, a peptide within H2 shows no differences between WT and A290P mutant (lower panel).

Because initial attempts to purify sufficient quantities of a senescence-domain-only tether-construct for lipid transfer experiments were unsuccessful, we prepared intact protein but with an engineered protease cleavage site between the PH-like and senescence domains. We expressed this modified construct for CtSpartinL in Expi293 cells and isolated it via FLAG-affinity resin, then removed N-terminal portions through the PH-like domain by proteolytic cleavage. This senescence-domain-only construct (CtSparinL_SD_, residues 186-480) transferred lipids comparably to full-length CtSpartinL (residues 7-480), indicating that the senescence domain is sufficient for lipid binding and transfer (Fig. 2E).

### The senescence domain binds LDs directly

Spartin associates with LDs in vivo, and the association is mediated by an N-terminal fragment of the senescence domain comprising two predicted amphipathic helices (Chung et al., 2023). To further characterize the LD targeting function of the senescence domain and in particular whether spartin’s interaction is directly with lipids or mediated by an adaptor protein, we carried out membrane binding studies in vitro. We incubated purified spartin with artificial LDs or with liposome preparations and then assessed association by flotation assay (Fig. 2F). HsSpartin bound robustly to both artificial LDs as well as liposomes comprising methyl branched diphytanoyl (4-ME) phospholipids, which introduces packing defects to better mimic the LD surface, but not to liposomes lacking 4-ME lipids (Fig. 2F). Further, we found that a construct lacking the senescence domain (ΔSD) did not bind to either LDs or liposomes, whereas a construct lacking only helices 3 and 4 of the senescence domain (ΔH3H4) still associated (Fig. 2F). Thus, spartin can interact directly with LDs, most likely via helices 1 and 2 of its senescence domain as reported (7). Several well studied LD targeting motifs feature amphipathic helices, which associate with LDs by integrating their hydrophobic face into the LD glycerophospholipid monolayer (16), and an attractive hypothesis (7) is that helices 1 and 2 of the senescence domain might insert similarly. Note, however, that typical helical LD-targeting motifs are unstructured in solution in the absence of LDs or membranes (for example, (17)), whereas helices 1 and 2 of the senescence domain are structured (below). Plausibly, the senescence domain might adopt different conformations, depending on its function in LD binding or lipid transfer.

### Structural perturbations in the senescence domain correlate with defects in lipid binding and transport

The senescence domain has no homology with any structurally characterized protein, including any known lipid transporters, and AI-based algorithms (18, 19) do not confidently predict its fold, whether as a monomer or multimer. Indeed, AlphaFold2 (19) predictions for the senescence domains of HsSpartin, CeSpartin, and CtSpartinL differ dramatically despite moderate sequence identities in the domain (24% between HsSpartin and CeSpartin, 28% between HsSpartin and CtSpartinL), suggesting the predictions are unreliable. Thus, for insights as to how this region of spartin solubilizes lipids to effect their transfer, and because we were unable to crystallize any senescence-domain-containing construct, we turned to hydrogen/deuterium exchange mass spectrometry (HDX-MS) (20, 21). This technique measures the exchange rate of amide hydrogens with deuterated solvent, where the primary determinant of exchange is the stability of secondary structure, allowing for assessment of secondary structure dynamics. Intrinsically disordered regions lacking secondary structure exchange very rapidly (on the order of milliseconds to seconds), whereas exchange is slower in secondary structure elements that form in folded protein. We used either CeSpartin or CtSpartinL in these experiments as yields of HsSpartin were not sufficient for HDX-MS analysis. For both CeSpartin and CtSpartinL we found that predicted helices 1-3 in the senescence region exhibit exchange consistent with the formation of secondary structure and folding (Fig. 2G). We also compared CtSpartinL with the lipid transport impaired mutant (A290P in helix 1), finding significant differences in hydrogen-deuterium exchange rates around helix 1 of the senescence domain (Fig. 2G-H), whereas other regions of the WT and mutant proteins behaved similarly. The HDX-MS data thus indicate that the senescence domain forms tertiary structure, making plausible that the senescence domain folds into a lipid transfer module, and that mutations leading to structural changes in the senescence domain impact lipid binding and transfer. Because spartin co-purifies with lipids (above), the HDX-MS analysis likely was carried out on the lipid-protein complex rather than apo-protein. We speculate that folding of the senescence domain may in fact require the presence of lipids, explaining the failure of current AI-algorithms to predict the tertiary structure.

### Loss of HsSpartin’s lipid transfer ability leads to defects in LD turnover

We next asked whether HsSpartin’s lipid transfer ability is relevant for LD turnover in cells. For functional studies, we used CRISPR-Cas9 technology to make a HsSpartin KO HeLa cell line (Fig. 3A). We induced LD formation in these or WT cells by treating them with oleic acid, then quantitated LD area and number and cellular TAG content 24 hours later. As in previous studies in HEK293T or SUM159 cells (6, 7), HeLa cells lacking spartin had more LDs (Fig. 3A) and a higher TAG content versus wild-type cells (Fig. 3B), consistent with a role for spartin in LD turnover. Neither spartin overexpression in WT cells nor short term (30 min) oleic acid treatment in spartin-KO cells affected cellular TAG content (Fig. 3B), in agreement with previous reports that spartin does not play a role in TAG synthesis (7). Consistent with the report that spartin delivers LDs to autophagosomes (7), we observed fewer LD-autophagosome contacts in KO cells than in cells stably expressing a WT construct (Fig. 3C); and in agreement with the proposed role for spartin in macrolipophagy (7), where lipid droplets are captured in autophagosomes and later internalized into lysosomes for degradation, we found reduced LD co-localization with and internalization into lysosomes in KO cells as compared to WT cells (Fig. 3D-E).

**Figure 3:**
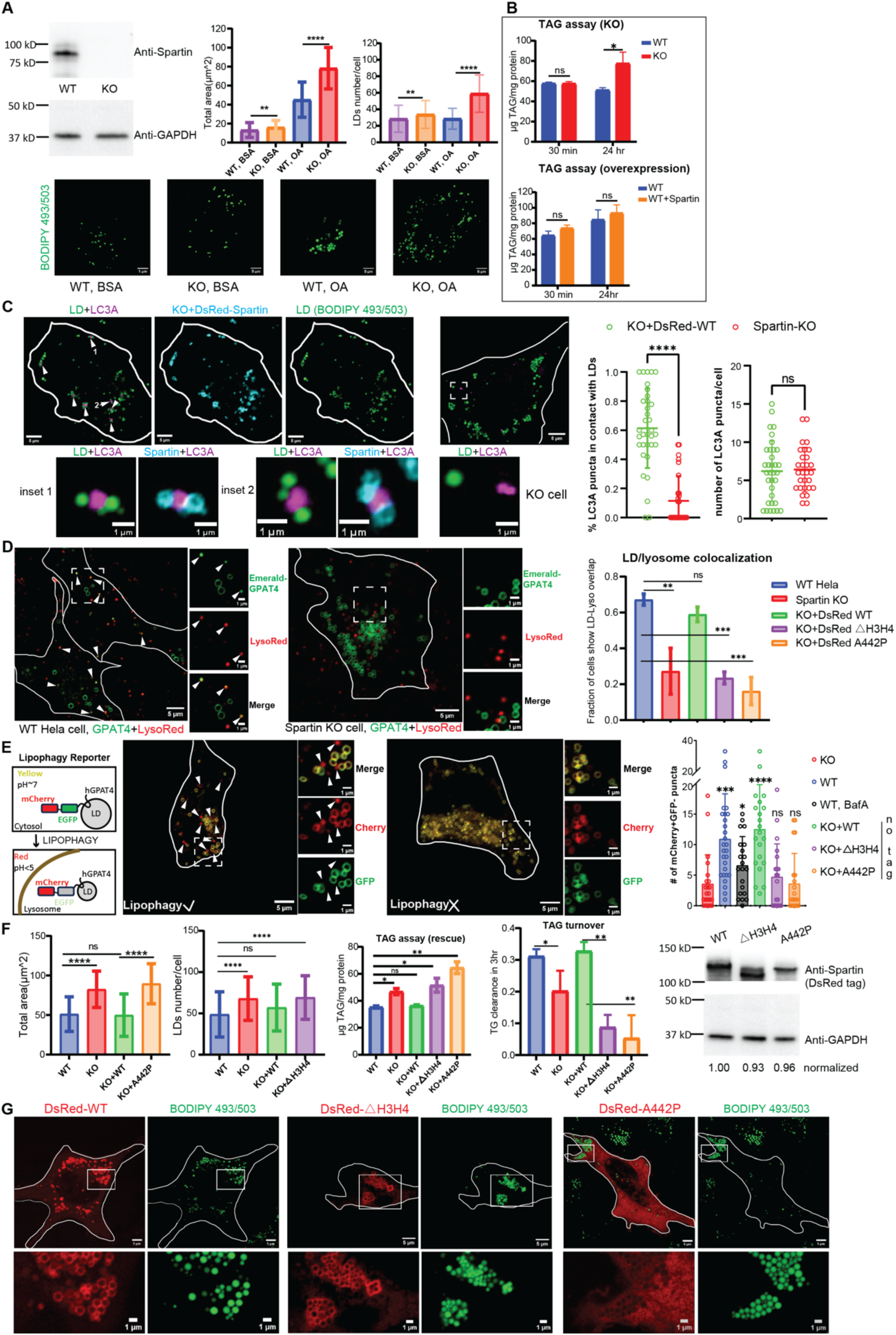
Loss of spartin’s lipid transfer activity leads to defects in LD turnover. **A**: Spartin knock out in Hela cells results in LD accumulation. Spartin was successfully deleted as in anti-spartin western blot. Spartin KO cells had significant LDs accumulation under basal condition (BSA treatment) and OA challenge (0.1 mM oleic acid for 18-24 hrs). Bar graphs: total area of LDs and number of LDs per cell (>100 cells per sample); fluorescence images in bottom row: BODIPY 493/503 staining of LDs in these cells. **B:** Spartin KO cells show TAG accumulation. TAG content in spartin KO cells was significantly higher than WT cells after 24-hr oleic acid treatment but was not altered after 30-min treatment (n=3 for each condition). Overexpression of spartin in WT Hela cells did not change TAG content 30 min or 24 hrs after 0.1 mM oleic acid treatment. **C**: Spartin brings LDs to LC3A autophagic membranes. BFP-LC3A construct was transiently expressed in spartin KO cells or cells stably expressing DsRed-Spartin WT construct (KO+DsRed-WT). When LC3A-LD-Spartin are co-imaged, a major portion of LC3A punctae (magenta) closely engage with spartin (cyan)-decorated LDs (green) and are highlighted by white arrows. The inset shows an enlarged view of these LC3A punctae. Quantification: percentage of LC3A punctae in contact with LDs was significantly smaller in spartin KO cells (n=29, 11%) versus in cells stably expressing a spartin construct (n=34, 61%), although the total number of LC3A punctae in cells was not affected. **D:** LD-lysosome colocalization depends on spartin’s lipid transfer activity. In cells (WT Hela, Spartin KO, KO+Dsred-WT, KO+Dsred-△H3H4 and KO+Dsred-A442P), a marker for LDs (mEmerald-GPAT4) was transiently transfected and lysosomes were stained by Lysotracker DeepRed. Fluorescence images show co-localization between Emerald-GPAT4 and lysosomes in WT Hela cells, but not in Spartin KO cells. A portion of LDs co-localize so exactly with lysosomes (indicated by white arrows) as to suggest internalization into lysosomes. Fraction of cells showing LD-lysosome co-localization/internalization was quantified for each sample; n>30 for each sample. **E**: Lipophagy is not rescued by lipid transfer deficient Spartin constructs. An environmentally sensitive probe, mCherry-EGFP-GPAT4 (25), was transfected into cells for monitoring LD internalization into lysosomes in the presence (left) or absence of spartin (KO cell, right). mCherry^+^GFP^-^ puncta were counted in each sample, n>20 for each cell type and statistical analysis was performed against KO cell. As a control for the probe, BafilomycinA (BafA, 100 nM) prevents lysosome acidification, reducing number of LDs visualized within the lysosome in WT cells. See Supplementary Fig. S1B for protein expression verification. **F:** WT spartin but not △H3H4/A442P constructs rescue the LD/TAG accumulation phenotype in spartin KO cells. Rescue experiments were performed in stable cell lines to quantitate LD abundance (n>70 cells per sample) and TAG content. To measure TAG content, cells were treated with OA for 24 hrs, then OA was removed. TAG levels were determined immediately after and 3 hours after OA removal to assess turnover; n=3. Western blotting indicates successful generation of stable cell lines; expression levels were normalized to KO+DsRed-WT construct. **G:** HsSpartin_ΔH3H4_ localizes to LDs like the spartin WT protein, but the A442P mutant is mis-localized. DsRed-Spartin constructs were transiently expressed in Hela cells, with 0.1 mM OA treatment overnight. Both WT Spartin (n=22) and ΔH3H4 construct (n=16) formed donut-shaped rings around LDs; in contrast the A442P mutant (n=16) is cytosolic and excluded from LDs in some cases. The same localizations were observed in cell lines stably expressing DsRed-Spartin constructs (data not shown). Scale bar, 1μm for inset. See Supplementary Figure S1C, showing that △H3H4 but not A442P mutant tethers LDs to LC3A-positive punctae.

LD turnover defects in the KO cells, as assessed by the total area of LDs per cell and TAG levels, could be rescued by expression of a full-length spartin construct but not by lipid-transport impaired senescence domain mutants, HsSpartin_A442P_ and HsSpartin_ΔH3H4_ (Fig. 3F). Lipophagy, as assessed by the number of lysosome-co-localized or -internalized LDs, was likewise rescued in KO cells by expression of the full-length spartin construct but not of transport-impaired mutants (Fig. D-E). Loss of function of HsSpartin_A442P_ most likely results from mislocalization and/or a loss of tethering versus impaired lipid transport ability as HsSpartin_A442P_ does not localize to LDs like WT constructs (Fig.3G). In contrast, HsSpartin_ΔH3H4_ localizes both to LDs and to LC3-positive autophagosomes like WT (Fig. 3G, Supplementary S1C). This finding indicates that HsSpartin_ΔH3H4’_s tethering capabilities are intact and implies that the senescence domain has an additional role beyond localization and tethering. Based on our in vitro data, we propose that this additional function is lipid transport.

### Possible mechanisms for spartin function

That senescence-domain containing HsSpartin, CeSpartin, and CtSpartinL, from three different species, all bind and transfer lipids in vitro bestows confidence that the senescence domain has lipid transport abilities. Further, our functional experiments with HsSpartin support that its lipid binding/transport ability is relevant for LD turnover and that it supports lipophagy (7). In lipophagy, LDs are fragmented before their uptake into autophagosomes (macrolipophagy) or lysosomes (microlipophagy) in a process poorly characterized in terms of lipid dynamics (2, 22). Fragmentation may be piecemeal (23, 24), involving the extrusion of small fragments from the LD that are subsequently engulfed by either the autophagosome or lysosome, or it may take place via lipolysis. Both LD deformation as in the piecemeal model or lipolytic breakdown into smaller droplets, which have a larger surface to volume ratio, may require augmentation of the glycerophospholipid monolayer around the neutral core of the LD. Thus, our results suggest the possibility that spartin could mediate glycerophospholipid transfer to LDs from a nearby organelle, perhaps the autophagosome or nearby ER, in order to grow the monolayer prior to breakdown. However, it is equally possible that LD breakdown before lipophagy involves removal of both glycerolipids and glycerophospholipids, and we do not exclude the possibility that spartin could transfer glycerophospholipids from the LD. Moreover, while our data indicate a role for spartin in lipophagy, they do not preclude that spartin might have a role in lipophagy-independent LD breakdown. Whatever the case, to our knowledge, our results are the first indication that lipid transfer proteins participate in LD turnover, including by lipophagy. Moreover, while we have investigated spartin only in the context of LD turnover, its lipid binding/transport ability could also feature in other still undiscovered aspects of LD biology and membrane homeostasis, and the role of lipid transport by spartin-like proteins is entirely unexplored. Precisely how spartin is regulated in the course of LD turnover and the role of spartin and spartin-like proteins in other physiological processes remain intriguing questions for future investigation.

## Materials and methods

### Materials

Rabbit polyclonal anti-*Hs*Spartin antibody (13791-1-AP) was purchased from Proteintech. Rabbit Anti-GAPDH antibody was from sigma (G9545). Lipids used for in vitro biochemical assay were purchased from Avanti Polar Lipids: DOPC (850375), DOPE (850725), Liver PE (840026), DGS-NTA (Ni; 709404), Liss-Rhod PE (810150), Brain PI (4,5) P2 (840046), NBD-PA (810173: 1-palmitoyl-2-{6-[(7-nitro-2-1,3-benzoxadiazol-4-yl)amino]hexanoyl}-sn-glycero-3-phosphate), NBD-PS (810192: 1-palmitoyl-2-{6-[(7-nitro-2-1,3-benzoxadiazol-4-yl)amino]hexanoyl}-sn-glycero-3-phosphoserine), NBD-PC (810130: 1-palmitoyl-2-{6-[(7-nitro-2-1,3-benzoxadiazol-4-yl)amino]hexanoyl}-sn-glycero-3-phosphocholine), NBD-PE (810153: 1-palmitoyl-2-{6-[(7-nitro-2-1,3-benzoxadiazol-4-yl)amino]hexanoyl}-sn-glycero-3-phosphoethanolamine), 18:0-18:1 PS (840039), 4ME-PC(850356), 4ME-PE(850402). BSA-OA complex (29557) and BSA control (29556) were purchased from Cayman chemicals. Sodium dithionite was from Sigma-Aldrich (157953). The Hela cell line (ATCC #CCL-2; RRID: CVCL 0030) was a gift from Mals Mariappan (Yale University, New Haven, CT), C. elegans cDNA was a gift from Daniel Colon-Ramos (Yale University, New Haven, CT). Plasmid for the 6xhis-PH-tethering construct (Bian et al., 2018) was a gift from Pietro De Camilli (Yale University, New Haven, CT). mEmerald-GPAT4 (152-208) was a gift from Jennifer Lippincott-Schwartz (HHMI, addgene plasmid #134468).

### Plasmid construction

*Hs*Spartin (Uniprot Q8N0X7) gene was synthesized by Genescript and cloned into pCMV10 plasmid with N terminal 3xFLAG tag for biochemical experiments, into pCMV10 plasmid with N-terminal 3xFLAG-DsRed tag for imaging and cellular experiments (transient transfection), or into pLVX plasmid with N-terminal DsRed tag for generating stable cell lines (lentiviral transduction). Point mutation or truncation constructs were generated using NEB Q5 site mutagenesis strategy (#E0554S). All constructs used for lipid transfer assay have a 25GS-6xHIS linker sequence as previously described (26).

The coding sequence for full length *Ce*Spartin (Uniprot O44735) was amplified from *C. elegans* cDNA library and subcloned into pET28-6xHIS-MBP-TEV digestion site plasmid. The sequence coding for residues 95-240 of *Ce*Spartin fragment was cloned into pET29 vector with C terminal 6xHIS tag for bacterial expression. The sequence coding for full length CeSpartin and residues 121-476 was cloned into pCMV10 plasmid with N-terminal 3xFLAG tag for Expi293 expression. Constructs used for lipid transfer assay have a 25GS-6xHIS linker sequence.

The sequence coding for residues 7-480 of CtSpartinL (Uniprot G0S883) was amplified from *C. Thermophilum* genomic DNA and cloned into pCMV10 plasmid with N terminal 3xFLAG tag for mammalian cell expression or pET28-6xHIS-MBP plasmid for bacterial expression. For obtaining a senescence-domain-only construct, a prescission protease cleavage site was engineered between A185 and D186 in the backbone of 3xFLAG-(7-480)-25GS-6HIS pCMV10 construct using NEB Q5 mutagenesis strategy. Point mutant and truncated constructs of *Ct*SpartinL protein were generated using NEB Q5 mutagenesis strategy. Construct used for lipid transfer assay have a 25GS-6xHIS linker sequence.

*Hs*LC3A (Uniprot Q9H492) gene was subcloned from a pGEX-GST-LC3A template into pCMV10 plasmid with N-terminal BFP tag for imaging experiment. The mCherry-EGFP-GPAT4 (152-208) construct was created using multi-fragment Gibson assembly strategy (NEB, E5510S).

Primers used in this study are in Table S1.

### Protein expression and purification

From *E. coli:* The full length CeSpartin construct (pET28-6xhis-MBP-TEV cleavage site-*Ce*Spartin) was expressed in Nico21 cells (C2529H, New England Biolabs). Cells were grown at 37°C to an OD600 of 0.6–0.8, when protein expression was induced with 0.3 mM IPTG, and then further cultured at 18°C overnight. Cells were pelleted, resuspended in buffer A (50 mM HEPES, pH 8.0, 500 mM NaCl, 1 mM TCEP, and 10% glycerol) containing 1× complete EDTA-free protease inhibitor cocktail (1187358001; Roche) and lysed in an Emulsiflex-C5 cell disruptor (Avestin). Cell lysates were clarified via centrifugation at 15000rpm (JA-20) for 30 min. To enrich CeSpartin, supernatant was incubated with Ni-NTA resin (#30210; Qiagen) for 1 h at 4°C in the presence of 20 mM imidazole, and then the resin was washed with buffer B (buffer A + 20 mM imidazole) for at least 4 column volume (CV). Retained protein was eluted from the resin with buffer A supplemented with 200 mM imidazole. TEV protease was added to digest overnight at room temperature. Post digestion, cleaved protein product was passed through Ni-NTA resin and amylose resin (E8021, NEB) to remove the TEV protease and HIS-MBP tag. The flow-through was collected, concentrated in a 30-kD molecular weight cutoff Amicon centrifugal filtration device (Sigma, UFC8030), and loaded onto a Superdex 200 10/300 column (GE Healthcare) equilibrated with buffer A. Peak fractions containing pure *Ce*Spartin were recovered and concentrated for biochemical assays. *Ce*Spartin proteins used for HDX-MS analysis were prepared similarly, except pH 7.4 buffer was used. The pET28-6xhis-MBP-TEV cleavage site-*Ct*SpartinL (7-480) constructs were expressed and purified similarly as full-length *Ce*Spartin.

The pET29-*Ce*Spartin (95-240)-6xHIS construct was expressed in BL21 (DE3) *E. coli* cells (69450; Sigma-Aldrich). Cells were grown at 37°C to an OD600 of 0.6–0.8, when protein expression was induced with 0.5 mM IPTG, and then further cultured at 18°C overnight. Protein was purified using Ni-NTA resin (20 mM imidazole and 40 mM imidazole for wash, 200 mM imidazole for elution) and loaded onto a Superdex 200 10/300 column (GE Healthcare) for analysis with buffer A. Peak fractions containing CeSpartin were used in biochemical assays.

The 6xhis-PH-tether construct was purified as described before (12). The E-Syt2 construct was purified as previously (11).

From Expi293 cells: Constructs in pCMV10 vector with N terminal 3xFLAG tag were transfected into Expi293F cells (A14527; Thermo Fisher Scientific) with PEI (Polyscience, 23966) for 48 hours according to the manufacturer instructions. The cell pellet was resuspended in buffer A supplemented with 1× complete EDTA-free protease inhibitor cocktail. Cells were lysed by 3-5 freeze-thaw cycles and centrifuged at 27,000 g for 30 min to clear the lysate. Lysates were incubated with M2 anti-FLAG resin (A2220; Sigma-Aldrich) for two hours at 4°C to capture FLAG-tagged protein. Flag resin was washed with buffer A and further incubated with buffer A containing 2.5mM ATP and 5 mM Mgcl2 overnight to remove chaperones. The protein was eluted with buffer A supplemented with 0.25 mg/ml 3× FLAG peptide (A6001; APExBio) for biochemical assays. *Ce*Spartin proteins used for lipidomic MS analysis were prepared similarly, except for extensive washes with buffer A (4 column volumes, or 60 ml, for 1-hr) to remove non-specifically bound lipids before elution. CtSpartinL proteins (WT and A290P) used for HDX-MS analysis were prepared similarly, except pH7.0 buffer was used, and proteins were gel filtrated on Superdex S200 10/300 and concentrated to 0.7 mg/ml for sample preparation.

The senescence-domain-only construct of CtSpartinL (3XFLAG-(7-480_Precission_)-25GS-6HIS) was expressed in Expi293 cells and purified via anti-FLAG resin as before, then incubated with Precission protease (Cytiva, 27084301) at 4C for overnight. After protease cleavage, flow-through was collected and incubated with TALON resin (TaKaRa, 635502) for 30 minutes at 4°C to capture the protein fragment comprising the senescence domain-25GS-6HIS, in the presence of 10 mM imidazole. TALON resin was washed three times with buffer A containing 20 mM imidazole and protein was eluted using 200 mM imidazole. Eluate was used in lipid binding assays or else dialyzed against buffer A to remove imidazole for other biochemistry.

### Lipid co-migration assay

NBD-labeled lipids in chloroform were dried using a N_2_ stream and resuspended in methanol at 1 mg/ml. 1ul of NBD-phospholipid substrates/methanol were incubated with purified proteins on ice for 2h. Samples were loaded onto 4–15% Mini-Protean Precast Native gels and run for 90 min at 100 V. NBD fluorescence was visualized using an ImageQuant LAS4000 (GE Healthcare). Then gels were stained with Coomassie blue G250 to visualize total protein. Images were analyzed via Fiji (https://fiji.sc).

### Liposome preparation

Lipids in chloroform were mixed (for donor liposomes: 61% 1,2-Dioleoyl-sn-glycero-3-phosphocholine [DOPC], 30% liver PE, 2% NBD-labeled lipids, 2% Rh-PE, and 5% DGS-NTA (Ni); for acceptor liposomes: 65% DOPC, 30% liver PE, and 5% PI[4,5]P2) and dried to thin films under vacuum for 30 min. Lipids were subsequently dissolved in assay buffer C (20 mM HEPES, 200 mM NaCl, 1 mM TCEP, pH8.0) at a total lipid concentration of 1 mM and incubated at 37°C for 1 h, vertexing every 30 min. Liposomes were subjected to 10 freeze–thaw cycles. Crude liposomes were then extruded through a polycarbonate filter with 100 nm pore size a total of 11 times via a mini extruder (Avanti Polar Lipids) and used within 24 h or kept at -80C until use.

### Artificial LD preparation

Artificial lipid droplets were prepared according to (14) with minor modifications. Briefly, as in (3), 2 mg of total phospholipids (61% DOPC, 30% DOPE, 2% NBD-labeled lipids, 2% Rh-PE, and 5% DGS-NTA by molar ratio) were mixed and dried to thin films. 5 mg of TAG (T7140; Sigma-Aldrich) was added to the phospholipid film and dried using a N_2_ stream for 10 min. The lipid mixture was further dried under vacuum for 30min, and then lipids were resuspended in 100 μl buffer C. The sample was vortexed for a total of 8 min, with 10s +/- cycle. The milky lipid mixture was centrifuged at 20,000 g for 5 min at 4C. The fraction containing lipid droplets floated as a pink band at the top of the tube. The underlying solution and pellet were removed. This process was repeated until no pellet formed upon centrifugation. The white band was resuspended in 100 μl buffer C and centrifuged at 1000 g for 5 min at 4C. The solution underneath the floating pink band was collected. Low-speed centrifugation was repeated until no pink band formed after centrifugation; the final preparation of artificial LDs was analyzed by positive stain TEM as in (14)(Supplementary Fig. S1A). The concentration of NBD-lipids in the final solution was determined by NBD fluorescence using liposomes containing the same ratio of NBD- and Rh-lipids as standards.

### In vitro lipid transfer assay

Lipid transfer experiments were set up at 30°C in 96-well plates, in 100 μl reaction volumes containing donor liposomes and acceptor liposomes. Proteins (at 0.25 μM, at ratios of 1 protein per 800 lipids) were added to start the reaction after 5-min pre-reading, and NBD emission (538 nm, excitation at 460 nm) was monitored for indicated time using a Synergy H1 Multi-Mode Microplate Reader (Agilent). For transfer from artificial LDs to liposomes, 80 μM lipids in donor LDs and 80 μM lipids in acceptor liposomes, and 0.1 μM proteins were used. Lipid transfer assays were performed similarly for the dithionite assay that controls for fusion (13), except for the addition of freshly prepared dithionite (5 mM final concentration) after the last reading point, and NBD fluorescence was monitored for an additional 5 min. As originally described, the dithionite fusion assay (13) used headgroup-incorporated NBD, whereas the NBD in lipids used for the transfer assays was incorporated in the acyl-chain. The assay also works for these lipids, as acyl-chain incorporated NBD in outer leaflet lipids is still accessible to the added dithionite (27).

### In vitro flotation assay

Proteins were purified from Expi293 cells as described above, and liposomes and artificial LDs were prepared as above, the phospholipid composition is 64.5% DOPC, 30% DOPE, 0.5% Rho-PE (for visualization after flotating), 5% 18:0-18:1 PS. For 4ME liposomes, 4ME-16:0 PC & PE were used. 50 μl of proteins were incubated with 50 μl of liposomes/artificial LDs at RT for 30 minutes. Post incubation, protein-liposome or protein-LD mixtures were combined with equal volume of 60% Optiprep solution (D1556-250ML, sigma) and placed at the bottom of the centrifuge tube (Beckman, 344090), and 10% Opti-prep and 0% Optiprep solutions were carefully pipetted on the top. After 1-hr of ultracentrifugation (SW55 rotor, 30k rpm, 4C), the tubes were immediately flash-frozen by LN2 and fractions were collected for SDS-PAGE.

### Lipidomic MS analysis of *Ce*Spartin-associated lipids

Purified *Ce*Spartin protein from Expi293 cell was sent to Michigan State University’s Collaborative Mass Spectrometry Core for untargeted global lipidomic analysis. The sample was spiked with internal standards and calibration mixture. Lipids were extracted with methyl tert-butyl ether twice, and after drying, resuspended in isopropanol containing 0.01% butylated hydroxy toluene. The sample was resolved by Shimadzu Prominence high performance LC and lipid species were detected by a Thermo Fisher Scientific LTQ-Orbitrap Velos mass spectrometer in both positive and negative ionization modes. Lipid species were quantified based on internal standards and summed by lipid class.

### Blue native electrophoresis analysis of spartin proteins

Spartin proteins including HsSpartin, CeSpartin and CtSpartinL were expressed in Expi293 cells, and purified using buffer A (50 mM HEPES, 500 mM Nacl, pH8.0, 1 mM TCEP, 10% glycerol) using anti-FLAG resin. Proteins were quantified using BSA standards in SDS-PAGE. Proteins (1-2 μg) together with BSA (1 μg) were loaded onto native PAGE 4%-16% Bis-Tris gel (BN1002BOX, invitrogen) for electrophoresis at 4C using NativePAGE running buffer and NativePAGE Cathode buffer (contains 0.02% Coomassie G-250, BN2007, invitrogen). NativeMark unstained protein standards (LC0725, invitrogen) and NativePAGE sample buffer (BN2003, invitrogen) were used. Protein gels were fixed and destained according to manufacturer direction. Distinct from native gels in the lipid-co-migration assay (above), where samples migrate according to their charge/mass ratio, here the Coomassie dye in the cathode buffer coats protein sample, which consequently migrates according to MW (28).

### Hydrogen-deuterium exchange mass spectrometry

#### Sample preparation

HDX reactions examining *Ce*Spartin were carried out in 40 µl reactions containing 20 pmol of protein. Exchange was initiated by the addition of 30 µl of D_2_O buffer (20 mM HEPES pH 7.5, 100 mM NaCl) to 10 µl of protein (final D_2_O [c]of 70.7% [v/v]). Reactions proceeded for 3s, 30s, 300s, and 3000s at 20°C before being quenched with ice cold acidic quench buffer, resulting in a final concentration of 0.6M guanidine HCl and 0.9% formic acid post quench. HDX reactions comparing WT *Ct*SpartinL and A290P *Ct*SpartinL were carried out in 20 µl reactions containing 20 pmol of protein. Exchange was initiated by the addition of 18 µl of D_2_O buffer to 2 µl of protein (final D_2_O [C] of 84.9% [v/v]). Reactions proceeded for 0.3s (3s on ice), 3s, 30s, and 300s at 20°C before quenching as described above. All conditions and timepoints were created and run in triplicate.

Fully deuterated (Max-D) samples were created for each condition as in (29): *Ce*Spartin was resuspended to 2 µM and *Ct*SpartinL (WT and A290P) was resuspended to 10 µM in 7M guanidine HCl. After vortexing, samples were heated to 90°C for 5 minutes and cooled down to 20°C. To initiate the exchange reactions, 30 µl of D_2_O buffer was added to 10 µl of denatured *Ce*Spartin, and 18 µl of D_2_O buffer was added to 2 µl of denatured *Ct*SpartinL proteins. Exchange proceeded for 10 minutes at 50°C before samples were cooled to 20°C and then 0°C. Samples were quenched using ice cold acidic quench buffer as described above. All samples were flash frozen immediately after quenching and stored at -80°C until MS analysis.

#### Peptide digestion and MS analysis

Protein samples were rapidly thawed and injected onto an integrated fluidics system containing a HDx-3 PAL liquid handling robot and climate-controlled (2°C) chromatography system (LEAP Technologies), a Dionex Ultimate 3000 UHPLC system, and an Impact HD QTOF Mass spectrometer (Bruker), as described in (30). *Ce*Spartin samples were run over one immobilized pepsin column (Trajan; ProDx protease column, 2.1 mm x 30 mm PDX.PP01-F32) at 200 µL/min for 3 minutes at 10°C, and *Ct*SpartinL samples were run over two immobilized pepsin columns (Waters; Enzymate Protein Pepsin Column, 300Å, 5µm, 2.1 mm X 30 mm) at 200 µL/min for 3 minutes at 2°C. The resulting peptides were collected and desalted on a C18 trap column (Acquity UPLC BEH C18 1.7mm column (2.1 x 5 mm); Waters 186003975). The trap was subsequently eluted in line with an ACQUITY 1.7 μm particle, 100 × 1 mm^2^ C18 UPLC column (Waters), using a gradient of 3-35% B (Buffer A: 0.1% formic acid; Buffer B: 100% acetonitrile) over 11 minutes immediately followed by a gradient of 35-80% B over 5 minutes. Mass spectrometry experiments acquired over a mass range from 150 to 2200 m/z using an electrospray ionization source operated at a temperature of 200°C and a spray voltage of 4.5 kV. Peptides were identified from non-deuterated samples of *Ce*Spartin, WT *Ct*SpartinL, or A290P *Ct*SpartinL using data-dependent acquisition following tandem MS/MS experiments (0.5s precursor scan from 150-2000 m/z; twelve 0.25s fragment scans from 150-2000 m/z). *Ce*Spartin MS/MS datasets were analyzed using PEAKS7 (PEAKS), and peptide identification was carried out using a false discovery-based approach, with a threshold set to 1% using a database of purified proteins and known contaminants (Dobbs et al., 2020). The search parameters were set with a precursor tolerance of 20 ppm, fragment mass error 0.02 Da charge states from 1-8, leading to a selection criterion of peptides that had a -10logP score of 20.3.

WT and A290P *Ct*SpartinL MS/MS datasets were analyzed using FragPipe v18.0, and peptide identification was carried out by using a false discovery-based approach using a database of purified proteins and known contaminants (Kong et al., 2017),(Dobbs et al., 2020),(Leprevost et al., 2020). MSFragger was utilized, and the precursor mass tolerance error was set to -20 to 20ppm, and the fragment mass tolerance was set at 20ppm. Protein digestion was set as nonspecific, searching between lengths of 4 and 50 aa, with a mass range of 400 to 5000 Da.

HD-Examiner Software (Sierra Analytics) was used to automatically calculate the level of deuterium incorporation into each peptide. All peptides were manually inspected for correct charge state, correct retention time, appropriate selection of isotopic distribution, etc. Deuteration levels were calculated using the centroid of the experimental isotope clusters. Results are presented as levels of deuterium incorporation. Back exchange is controlled with a maximally deuterated sample for each condition.

In comparing WT CtSpartinL with A290P CtSpartinL, differences in exchange in a peptide were considered significant if they met all three of the following criteria: ≥4.5% change in exchange, ≥0.45 Da difference in exchange, and a p value <0.01 using a two tailed student t-test. These samples were only compared within a single experiment and were never compared to experiments completed at a different time, or with a different final D_2_O level. To allow for visualization of the overall deuteration levels, we utilized percent deuteration (%D) plots (Fig. 2G). These plots show the Max-D corrected percentage of deuterium incorporation at the3 second timepoint for each condition, with each point indicating a single peptide. The Max-D corrected deuterium incorporation percentage for peptides with significant differences are highlighted in Fig. 2G with red rectangles, with the raw data for all analyzed peptides in the source data.

The data analysis statistics for all HDX-MS experiment are in Table S2 according to the guidelines of (31). The mass spectrometry proteomics data have been deposited to the ProteomeXchange Consortium via the PRIDE partner repository (32) with the dataset identifier PXD043175.

### Functional experiments

#### Cell culture and transfection

Hela cells were cultured in DMEM (11965092; Thermo Fisher Scientific) supplemented with 10% FBS (10438062; Thermo Fisher Scientific) and 1% penicillin-streptomycin (15140122; Thermo Fisher Scientific) at 37°C in a 5% CO2 incubator. Plasmid transfection was performed using Lipofectamine 3000 (L3000015; Thermo Fisher Scientific) according to the manufacturer’s instruction, or FuGene HD transfection reagent (Promega, E2311) as noted.

#### Generation of human Spartin KO cell line

CRISPR gRNAs targeting the sixth exon of Spartin were designed using the online TKO CRISPR Design Tool (https://crispr.ccbr.utoronto.ca/crisprdb/public/library/TKOv3/). 5 ′ - ATAGCGAAGCAAGCTACAGG-3′ was chosen as gRNA to knock out Spartin. The gRNA was cloned into pX458 (48138; Addgene) as described previously (33). Hela cells were transiently transfected with the constructs containing gRNA. After 48 h, the GFP positive individual cells were selected by flow cytometry (BD FACSAria) and seeded in 96-well plates for single clones. The clones were validated by PCR genotyping and western blot. For genotyping, genomic DNAs from single clones were extracted using QuickExtract DNA extraction solution (QE0905T; Lucigen), and PCR products containing the site of Cas9 targeting site were generated using the following primers: forward primer: 5 ′ -TGCTTCCTGGGTGAGTTGGGGTT-3 ′ ; reverse primer: 5 ′ - CACACATTCTCCTCTCCAACACATCAG-3′. Spartin KO clone has nucleotide insertion that leads to premature stop codon upon PCR product sequencing.

#### Generation of stable cell lines expressing spartin constructs

DsRed-Spartin constructs were generated in pLVX plasmid. HEK293 cells were cultured in DMEM medium (without antibotics) at 50%-80% confluence for virus transduction. pLVX-DsRed-Spartin plasmid together with psPAX2 and VSV-G plasmid were transfected into HEK293 cells using Lipofectamine 3000 (L3000015; Thermo Fisher Scientific) for generating lentivirus. Virus was collected at 48 h and 72 h after transfection by centrifuging the culture medium at 1500 g and filtering the supernatant using 0.45 μm membrane (Sigma, SLHVR33RS). Virus-containing medium was added into Spartin KO Hela cells for viral transduction. Two-day post viral transduction, cell debris were washed away by PBS and cells were treated with 10 μg/ml puromycin (Invitrogen, ant-pr) for selection purpose. Puromycin containing medium was used until all control cells (Spartin-KO cell without viral transduction) were dead. Stable cells were then further cultured for Anti-Spartin western blot to examine protein expression levels before being used for functional experiments. Stable cells expressing untagged Spartin constructs were made the same way.

#### Western blotting

Samples were denatured by the addition of SDS sample buffer (125 mM Tris-HCl, pH 6.8, 16.7% glycerol, 3% SDS, 0.042% bromophenol blue) and heated at 95°C for 5 min. The proteins were separated using denaturing SDS-PAGE and analyzed by immunoblotting. The proteins were transferred onto a nitrocellulose membrane. Then the membranes were incubated in 5% nonfat milk–TBST (20 mM Tris-HCl, pH 7.6; 150 mM NaCl; 0.1% Tween 20) for 2-hr at room temperature/overnight at 4C. The membranes were then washed three times using TBST and incubated with primary antibodies at 1:4000 dilution in 5% BSA-TBST at room temperature for 90 min. The membranes were washed three times and incubated for 60 min with secondary antibodies (1:4000) coupled to horse-radish peroxidase in TBST containing 5% milk. The blot was developed by ECL (BioRad) and visualized using ImageQuant LAS4000 (GE Healthcare).

#### Lipid extraction and TAG measurement were carried out as in (3)

Cells in 10-cm dishes were treated with 0.1 mM oleic acid (OA) for indicated times. Cells were washed twice with PBS and collected in PBS with cell scrapers; 1/10 of the sample was set aside for protein quantification. Protein amount was determined by Bradford (Bio-Rad) assay at 595 nm absorbance. Lipid extraction was performed as in (34). Briefly, the cell pellet was resuspended in hexane-isopropanol (3:2) solvent, then agitated at room temperature for 30 min to extract lipids. The mixture was spun at 3000 rpm for 10 mins. The organic solvent was transferred into a glass tube and dried overnight in a chemical hood. The lipid film was resuspended in 200 μl methanol-chloroform (2:1) and vortexed. The samples and a series of TAG standard dilutions were mixed with enzyme solution in the TAG assay kit (10010303; Cayman) and incubated at 37°C for 30 min. Absorbance at 540 nm was monitored. To assess TAG clearance, OA was removed 24hrs after its addition, and TAG was measured immediately and after a further 3 hrs. TG clearance was calculated as (TAG_0hr_ – TAG_3hr_)/ TAG_0hr._

#### Live cell imaging. For LD abundance quantification

Cells were plated on glass-bottomed 35 mm Mattek dish and were treated with 0.1 mM OA (29557; CAYMAN) or same concentration of BSA control (29556; CAYMAN) for 18–24 hr if imaging LDs. Two-days after transfection, cells were incubated with BODIPY 493/503 (#D3922; Thermo Fisher Scientific) at 1 μg/ml for 30 min, washed with PBS, and subjected to 1× live cell imaging solution (A14291DJ; Thermo Fisher Scientific) for imaging High throughput confocal microscopy was performed on Nikon Ti2-E inverted microscope with a 63× oil-immersion objective, CSUX1 camera Photometrics Prime 95B, Agilent laser 488 nm, and a 37°C chamber. Images were acquired using Nikon Elements and analyzed in Fiji (ImageJ). Different samples were imaged in one session with the same settings. For LD analysis, maximum projection of Z-stack was performed, and the images were set at the same threshold. LDs area and number were analyzed after converting to binary format.

#### For LC3A-Spartin-LD imaging

Cells (Spartin KO Hela cell or Hela cell stably expressing DsRed-Spartin constructs) were plated on glass-bottomed 35 mm Mattek dishes and transfected with BFP-LC3A plasmid the next day using FuGene HD transfection reagent. Next day, 0.1 mM OA was supplied to stimulate LD production. 18 h after OA incubation, OA-containing medium was replaced with regular DMEM medium. Cells were cultured for 3hr more before staining lipid droplets with BODIPY 493/503 (as described above) and imaged with 455/488/561nm laser on Zeiss LSM 880 point scanning confocal microscope at Yale CCMI facility. Images were first processed using Airyscan Processing in Zen Black, and all images were analyzed in Fiji (ImageJ).

#### For mEmerald-GPAT4 and Lysosome imaging

Cells were plated on glass-bottomed 35 mm Mattek dishes and transfected with mEmerald-GPAT4 plasmid using FuGene HD transfection reagent. 24hr post transfection, 0.1 mM OA was supplied to stimulate LD production. 18 h after OA treatment, OA-containing medium was replaced with medium containing lipoprotein deficient serum (880100, KALEN Biomedical). Cells were cultured for 3hr more before staining lysosome with Lysotracker DeepRed (L12492, thermos fisher) at 50 nM for 30 mins and imaged with 488/633nm laser in Airyscan mode on Zeiss LSM880.

#### For mCherry-EGFP-GPAT4 imaging

Cells were plated on glass-bottomed 35 mm Mattek dishes and transfected with *mCherry-EGFP-GPAT4* plasmid using FuGene HD transfection reagent. 24hr post transfection, 0.1 mM OA was supplied to stimulate LD production. 18 h after OA treatment, OA-containing medium was replaced with medium containing lipoprotein deficient serum (880100, KALEN Biomedical). Cells were cultured for 3hr more before imaging with 488/561nm laser in Airyscan mode on Zeiss LSM880.

#### Statistical analysis

Statistical analysis between groups was performed using Prism (GraphPad Software, RRID: SCR_002798; http://www.graphpad.com) with Welch’s two-tailed unpaired t test. Results were indicated in the following manner: * for P < 0.05, ** for P < 0.01, *** for P < 0.001, **** for P < 0.0001, where P < 0.05 is considered as significantly different.

## Contributions

NW and KMR conceived the project; NW designed the experiments and performed the biochemical assays and localization studies; NW and HZ generated the Hela cell lines and performed functional experiments together. MP and JEB carried out the HDX-MS analysis; JK helped with imaging strategy and artificial LD preparation; KMR supervised all aspects of this work with help from TJM for functional assays and JEB for HDX-MS experiments. KMR and NW wrote the manuscript, with inputs from TJM, MP and JEB.

## Acknowledgements

KMR thanks T. Levine for discussions that inspired this project, and we are grateful to T. Walther for discussion and to M. Henne for his constructive critique. KMR and TJM are supported by NIGMS (R35GM131715, R01135GM135290), JEB by NSERC (Discovery Grant 2020-04241) and salary awards from the Michael Smith Foundation for Health Research (Scholar 17686). NW and ZH received support from the China Scholarship Council.

---

**Figure S1.**
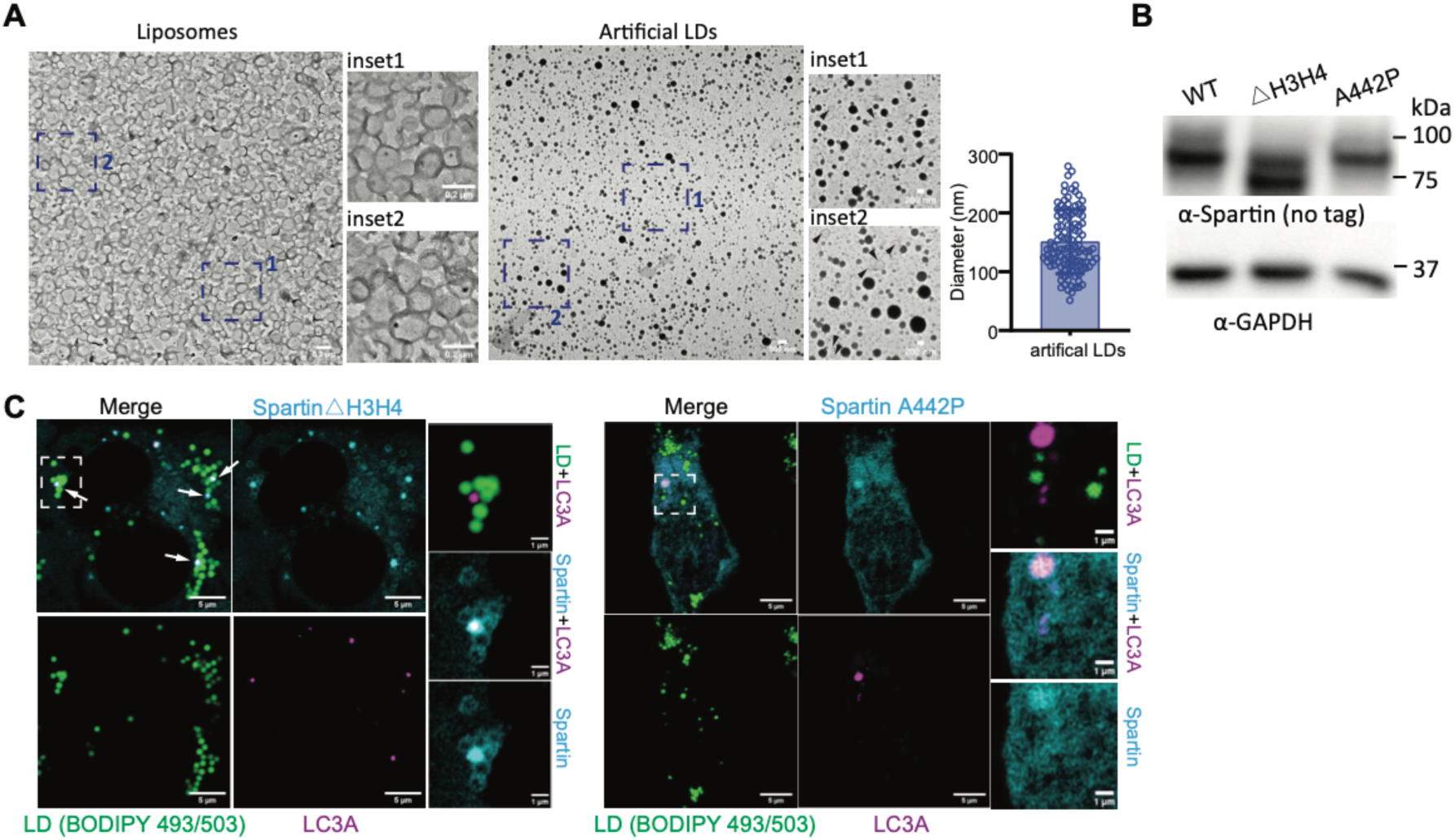
Examination of artificial LDs by EM and characterization of spartin mutants. **A**: Positive staining TEM images of liposomes and artificial LDs (used in Fig1H and Fig2F). Artificial LDs enriched with unsaturated TAG (average diameter=150nm, n>100) are stained by osmium tetroxide as electron-dense black dots. In contrast, liposomes are delineated by lower contrast grey membranes, and their interior appears white. In the insets for the lipid droplet preparations, arrows indicate liposome contaminants. Imaging was carried out using a T12 microscope with a magnification of 2700x at 120kV as in (14). **B**: Western blot shows successful generation of stable cell lines (used in Fig3E) expressing untagged Spartin constructs in KO background. **C**: Fluorescence microscopy shows Spartin_△H3H4_ maintains the ability to tether LD to LC3A-autophagosomes, but Spartin A442P mutant fails to tether as A442P does not localize to LDs anymore (Fig 3G). Stable cells expressing DsRed-Spartin △H3H4 (n=11) and DsRed-Spartin A442P construct (n=9) were transiently transfected with BFP-LC3A and followed by 0.1 mM OA treatment for 18-hr; OA was removed 3hrs before live cell imaging.

**Table S1.**
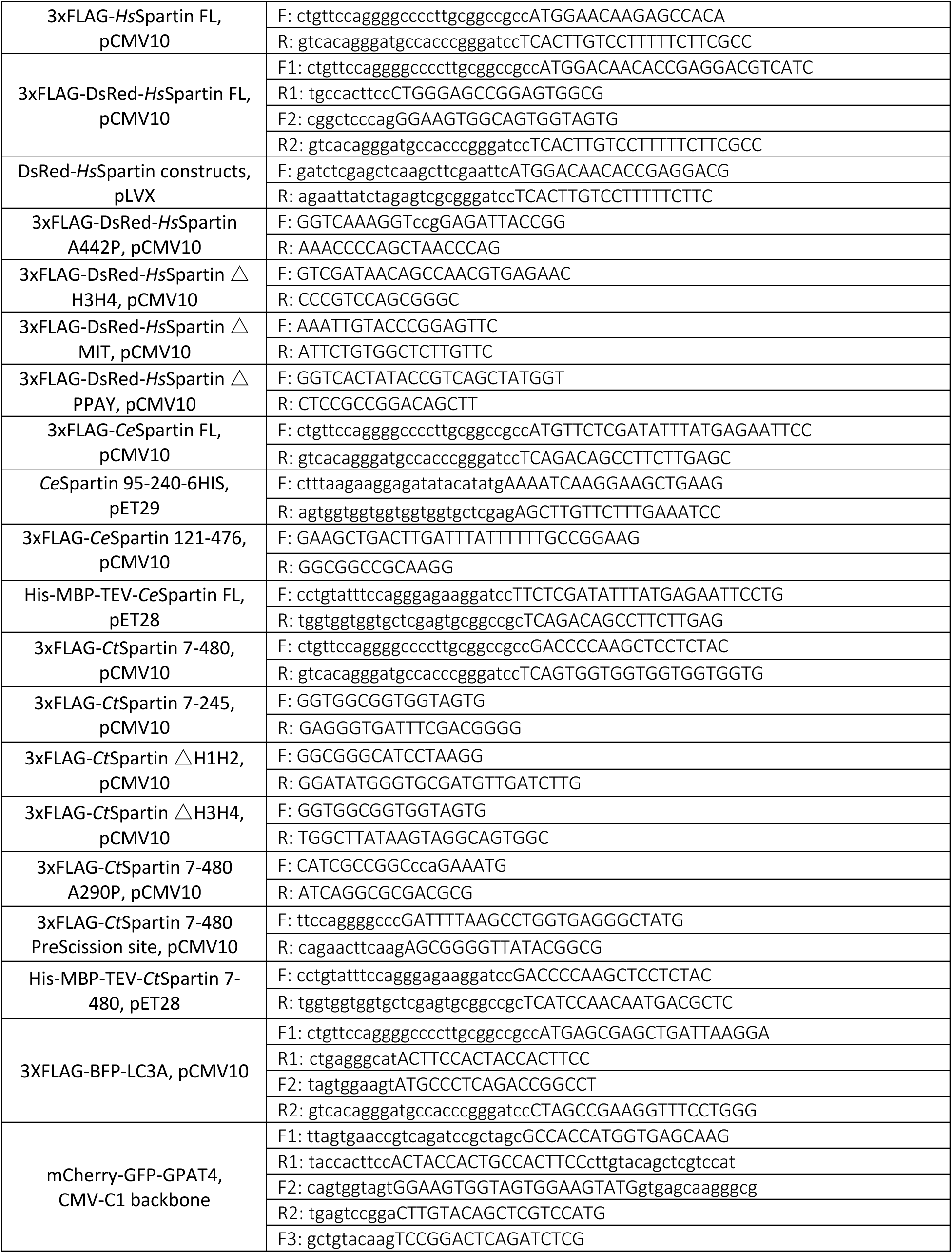
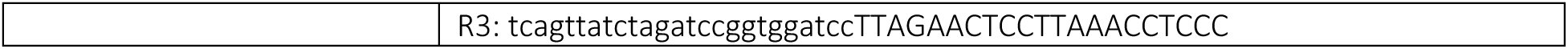

**Table S2.**

| Data set | CeSpartin FD-Corrected | CtSpartinL WT | CtSpartinL A290P |
| --- | --- | --- | --- |
| HDX reaction details | %D<sub>2</sub>O=70.7%<br>pH<sub>(read)</sub>=7.5<br>Temp=18°C | %D<sub>2</sub>O=84.9%<br>pH<sub>(read)</sub>=7.5<br>Temp=18°C | %D<sub>2</sub>O=84.9%<br>pH<sub>(read)</sub>=7.5<br>Temp=18°C |
| HDX time course (seconds) | 3s,30s,300s,3000s | 0.3s (3s on ice),<br>3s,30s,300s | 0.3s (3s on ice),<br>3s,30s,300s |
| HDX controls | Maximally deuterated sample | Maximally deuterated sample | Maximally deuterated sample |
| Back-exchange | 33.4% ± 16.0% | 33.3% ± 13.9% | 30.8% ± 13.6% |
| Number of peptides | 180 | 110 | 110 |
| Sequence coverage | 94.4% | 90.3% | 90.3% |
| Average peptide length/redundancy | Length= 14.8<br>Redundancy= 5.6 | Length= 17.5<br>Redundancy= 3.8 | Length= 17.5<br>Redundancy= 3.8 |
| Replicates | 3 | 3 | 3 |
| Repeatability | Average StDev=1.3% | Average StDev=1.1% | Average StDev=1.1% |
| Significant differences in HDX | >4.5% and >0.45 Da and unpaired t-test ≤0.01 | >4.5% and >0.45 Da and unpaired t-test ≤0.01 | >4.5% and >0.45 Da and unpaired t-test ≤0.01 |

